# Urbanisation, Bird Species Richness and Abundance Within Ibadan Metropolis, Nigeria

**DOI:** 10.1101/2021.05.18.444309

**Authors:** Festus O. Adegbola, Taiye Adeniyi Adeyanju, Soladoye B. Iwajomo, Ibukunoluwa Augustine Ayodele

## Abstract

Urbanisation is considered as one of the most profound threat to wildlife, with habitat loss and fragmentation being predominant. This study assessed the impacts of urbanisation on richness, abundance of bird species within Ibadan metropolis, Nigeria. A uniform grid of 500 square meters was installed on the map of Ibadan Metropolis using QGIS to produce 499 grid points distributed across the five urban local governments. 100 grids were selected randomly, identified with mapinR software application and surveyed with 5 point counts within each gird, established at 200m interval to avoid double counting. Each point count was observed for 5 minutes using a pair of 8×42mm binoculars within a 50m radius. Habitat variables like number of buildings, trees, paved roads, communication masts were also recorded.

A total number of 56 species of birds were observed at the end of the assessment, classified into 30 families. The test of statistics showed that there was no statistically significant difference in bird species richness between the Local Governments. The test of statistics showed that there was no statistically significant difference in bird species abundance between the local governments. The test of between-subjects effects revealed that there were no statistical significant effects when all the habitat variables were computed in the model on species richness. The number of paved roads and number of vehicles showed a significant effect on bird species abundance while others variables in the model did not exact statistically significant effects on bird species abundance.

The study therefore concluded that habitat actions due to urbanisation have not affected the richness and abundance of birds species found in all the local governments. The only habitat variables that have currently exacted significant effect on species abundance within the metropolis are number of vehicles and paved roads.

## INTRODUCTION

In wildlife conservation and management, there is need for a thorough understanding of the relationships among individual organisms and their environment, as it is important in the development of ecological theories and in the implementation of conservation strategies (Walker et al. 2008). It is general knowledge among environmentalists that there is an increase in the proportion of the earth’s surface now converted to human-dominated urban areas; it therefore becomes expedient to understand how urban wildlife communities within these now sophisticated ecosystem types are structured (McKinney 2002). The importance of understanding the ecological effects of urbanisation cannot be under-emphasised, especially with its rapid conversion of previously wild lands around the world. The process associated with urbanisation has profound effects on the distribution of wildlife species and their habitats (Wolff et al. 2018). Urban development is characterised by rapid population growth and profound land use transformation, leading to land conversion, which is a predominant process affecting ecological community structure and population dynamics of living organisms (Hostetler 1997). Research studies show that urban landscapes supports biotic communities in which only a few species increase in density compared to natural areas, resulting in a distinct difference the community diversity between these two landscapes (McKinney 2002).

Developing countries have a large number of wildlife existing outside the protected areas, on farmlands, and in urban areas. Among all wildlife, bird species are largely one of the most common wildlife surviving in urban communities (Gatesire et al. 2015). Birds are one of the most easily studied taxa occurring in cities worldwide as they serve as indicators variables in ecological assessment and monitoring (Magle et al. 2012). Birds are also important in maintaining ecosystems. For example, insectivorous bird species regulate disease vectors including mosquito and rodents. Pied Crow (*Corvus albus)*, contributes to biomass recycling and reduce levels of disposable waste as scavengers. Frugivorous birds play a crucial role in the dispersal of seeds of fruit trees. Sun birds also helps to pollinate plants (Gatesire et al. 2015).

Studies show that bird species in different regions respond to urbanisation in a similar way with most research suggesting that bird communities are negatively impacted by urbanisation (Lin et al. 2011; Sol et al. 2014). There is a general shrinkage in species distribution as urbanisation increases, and the fact that similar bird species can be found in various urban landscape indicates that urbanisation has a similar effect on local communities of birds irrespective of the region (Lin et al. 2011). As a result of the crucial role birds play in maintaining ecosystems and also support biodiversity, conservationists seek their protection against biological threats and protect the environment efficiently (Gatesire et al. 2015).

Primarily the development level, habitat diversity, age and diversity of vegetation present determine the richness of birds in urban areas (McKinney 2002). Native bird species that persists in urban landscapes partly depends on the actions of the landowners because the structural and vegetative characteristics of urban landscapes are largely human-influenced (McCaffrey and Mannan 2012). Urban areas have less assembly of bird species than adjacent natural areas, though some type and level of development support more native bird species than others. According to McKinney (2002), the moderate level of development and vegetation linkable to low density residential areas can support higher densities of some native birds species than other types of urban land use, also including undisturbed sites.

## MATERIALS AND METHODS

### STUDY AREA

The study was conducted in Ibadan metropolis, the capital of Oyo State. Ibadan metropolis, covering an area of 129.65 km^2^, is located in South-Western Nigeria in the southeastern part of Oyo State at about 119 kilometers (74 miles) northeast of Lagos and 120 kilometers (75 miles) east of the Nigerian international border with the Republic of Benin (Popoola and Wahab 2018). Ibadan falls totally within the forest zone but close to the boundary between the forest and the derived savanna. It lies between latitude 3?3′ N and 4?10′ N and longitude 7?2′E and 7?40′ E. (Popoola and Wahab 2018). The population of Metropolitan Ibadan is 1 338 659 according to census results for 2006 (Areola and Ikporukpo 2018). There are eleven (11) local governments in Ibadan Metropolitan area consisting of five urban local governments and six semi-urban local governments. The former are: Ibadan South East, Ibadan North East, Ibadan North West, Ibadan South West and Ibadan North Local Government Areas. Ibadan North Local Government has the largest land area among the urban Local Governments, while Ibadan North West is the smallest (Fig. 2). The second largest local government in Urban Ibadan is Ibadan South West. Ibadan metropolis is an important commercial centre and it comprises of people of different cultural and socio-economic backgrounds. General land use pattern of the Ibadan metropolitan area shows a clear distinction purely residential use for Urban Ibadan and agricultural use for semi-urban Ibadan (Salami et al. 2016).

**Figure 1:**
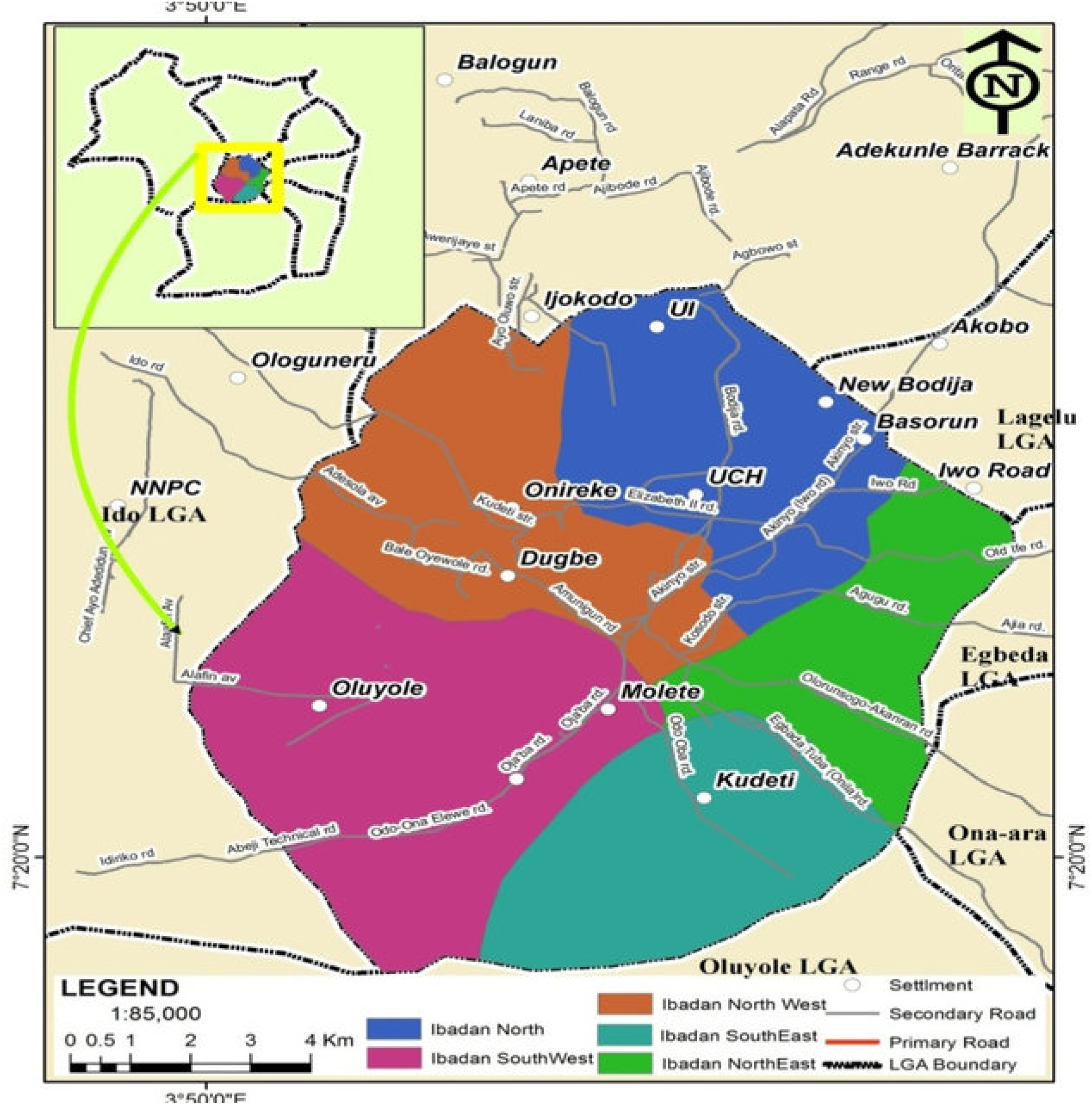
Map of Ibadan Metropolis Source: (Popoola and Wahab 2018)

**Figure 2:**
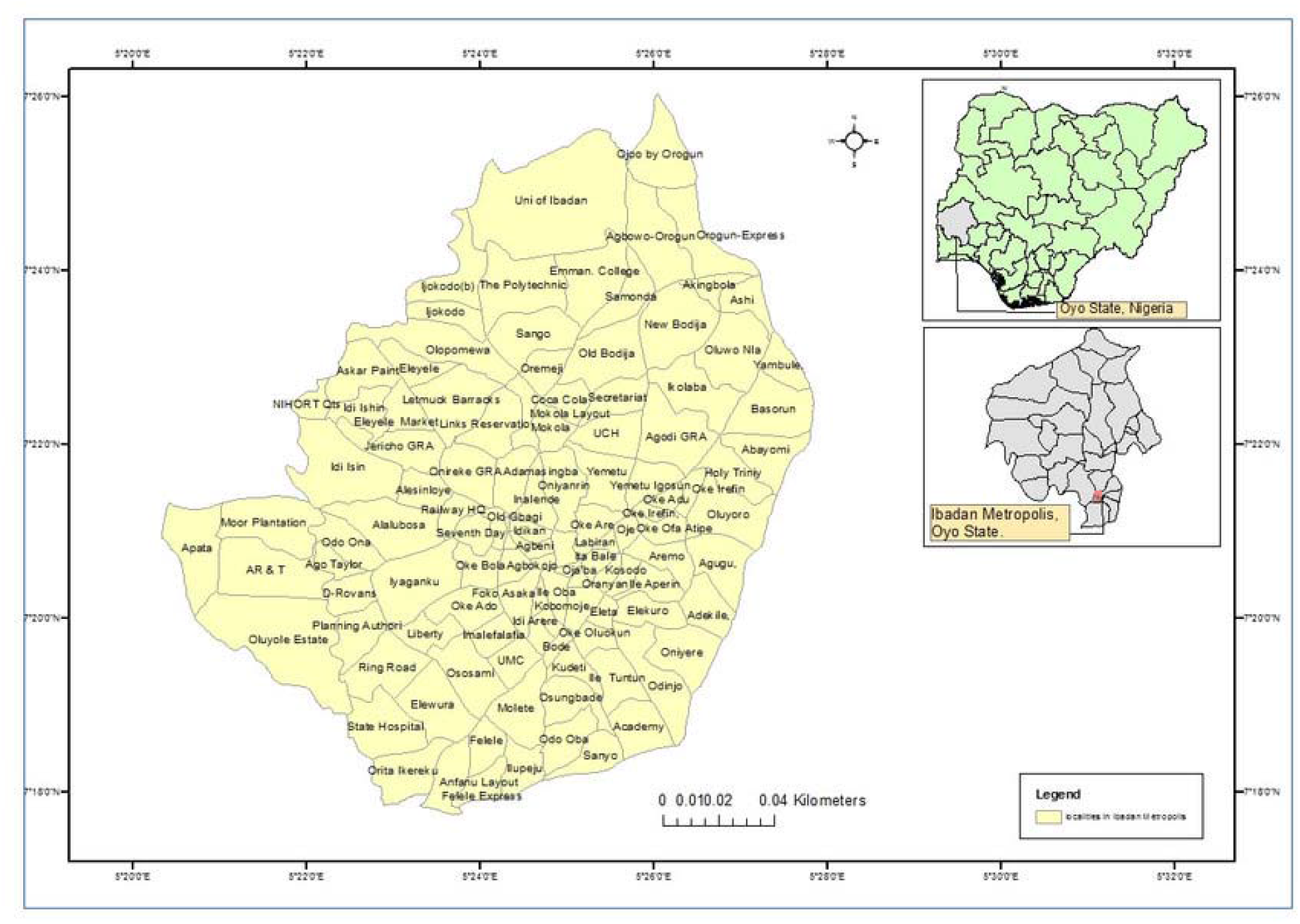
Map of Ibadan Metropolis, Oyo State showing communities Source: (Areola and Ikporukpo 2018)

### BIRD ASSESSMENT

Using QGIS software application, a uniform grid of 500 by 500 metres was installed on the map of Ibadan Metropolis to produce 499 grids distributed across all the five urban local governments. 100 grids were surveyed with 5 point counts within each gird, established at 200 m interval to avoid double counting, each point count was observed for 5 minutes within a 50 m radius. Laser range finder was used to delineate the 50 m radius around each survey point, and to estimate distances to birds. Birds encountered outside the study grids were recorded only when it has never been observed in any of the grids before. When a bird could not be identified in the field, photos from a high resolution camera were taken for later identification by an expert in ornithology. During each visit, a pair of 8×42 mm binoculars was used for sighting birds. Helms field guide to the birds of western Africa (Borrow and Demey 2013) was used to identify the birds; birds call were recorded with a voice recorder and later played back for confirmation. The data were collected between October to December 2020, and observations were done in the morning and evening. Habitat variables recorded for this study, the number of buildings, number of trees, number of communication masts, ground cover, canopy cover, number of pedestrians and vehicles passing within 25 m radius around the point, describe urban form. No specific permissions were required to conduct this work. All bird surveys were conducted in areas which are open to the public; therefore there was no need to ask land managers for approval

**Figure 3:**
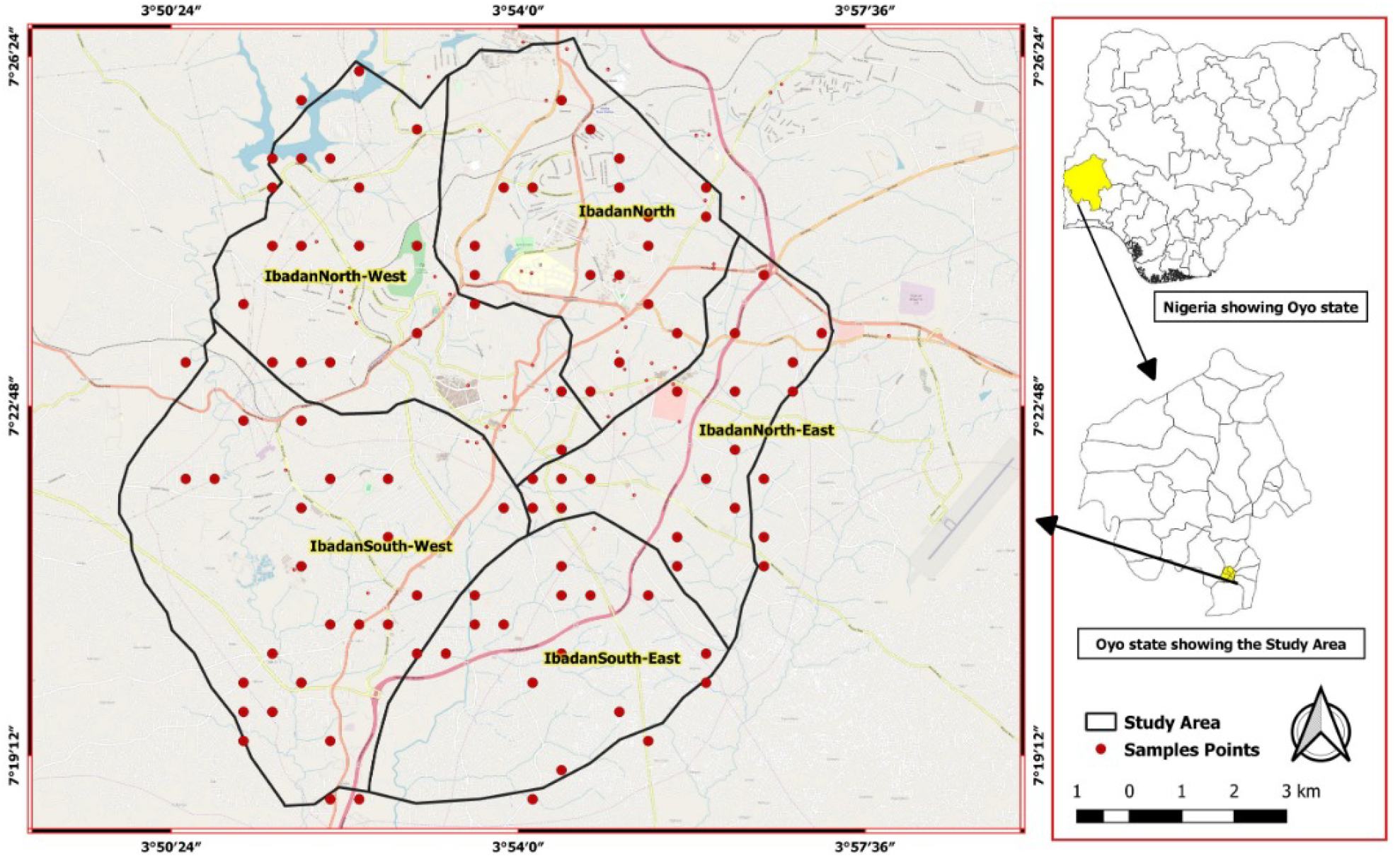
Map showing survey grids Source: QGIS Application

### DATA ANALYSIS

Species richness and abundance was obtained by counting the total number of species and individual per species respectively as recorded within each grid. The data collected were entered and summarised in Microsoft Excel spread sheet computer software for analysis. Tables and figures were used to represent the results.

General Linear models method was fitted in R statistical package version 3.4.2 using test of correlation to the determine relationship between the vegetation variables, bird species richness, and bird abundance at 0.05 level of significance.

## RESULTS

This chapter present the results of the data obtained for the purpose of this study. QGIS software application, using a uniform grid of 500 by 500 metres installed across all five selected urban local government in Ibadan was used in collecting the data. General Linear Model (GLM) of statistical analysis was used in analyzing the collected data and testing for statistical significant difference in bird species richness, bird abundance and the habitat variables that determines/ influence bird species richness and its abundance within the metropolis. (α = 0.05).

The study observed 56 different species of birds which was grouped into 30 families. (See appendix)

### DETERMINATION OF SPECIES RICHNESS WITHIN THE METROPOLIS

Descriptive statistics, Kruskal-Wallis H Test and Post Hoc Test were carried out to determine whether there are statistically significant differences between bird species richness and the five selected urban local government. The local government being the independent variable and bird species observed being the dependent variable.

The results showed that mean bird species richness at Ibadan North East, North West, South East and South West were estimated at 8.20 ± 1.643, 10.84 ± 2.630, 8.27 ± 2.724, 9.36 ± 2.629 and 10.54 ± 5.222 respectively.

On the overall, Bird species richness for Ibadan metropolis was estimated at 9.73 ± 3.301. (See table 1a)

**Table 1a:** Descriptive Statistics on Species Richness.

| **Local Government** | **N** | **Mean** | **Std. Deviation** |
| --- | --- | --- | --- |
| Ibadan North East | 5 | 8.20 | 1.643 |
| Ibadan North | 19 | 10.84 | 2.630 |
| Ibadan North West | 11 | 8.27 | 2.724 |
| Ibadan South East | 22 | 9.36 | 2.629 |
| Ibadan South West | 13 | 10.54 | 5.222 |
| Total | 70 | 9.73 | 3.301 |

**Table 1b:** Kruskal-Wallis H Test on species richness.

| Local Government | N | Mean Rank |
| --- | --- | --- |
| Ibadan North East | 5 | 24.70 |
| Ibadan North | 19 | 44.74 |
| Ibadan North West | 11 | 25.32 |
| Ibadan South East | 22 | 34.55 |
| Ibadan South West | 13 | 36.38 |
| Total | 70 |  |

| Test Statistics |  |
| --- | --- |
| Species richness |  |
| Chi-Square | 8.253 |
| Df | 4 |
| Asymp. Sig. | .083 |

a. Kruskal Wallis Test
b. grouping variable: local government

**Table 1c:**
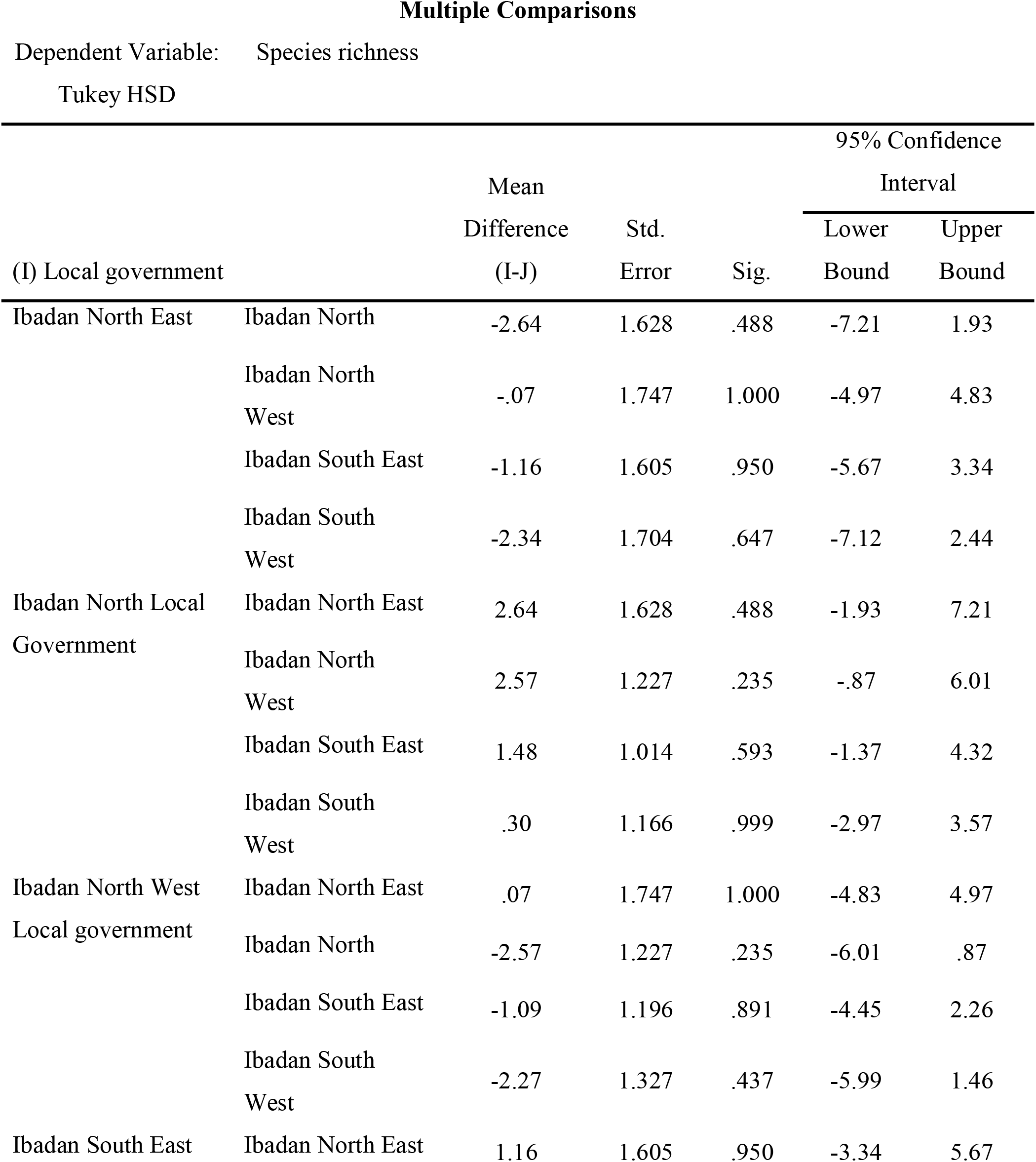
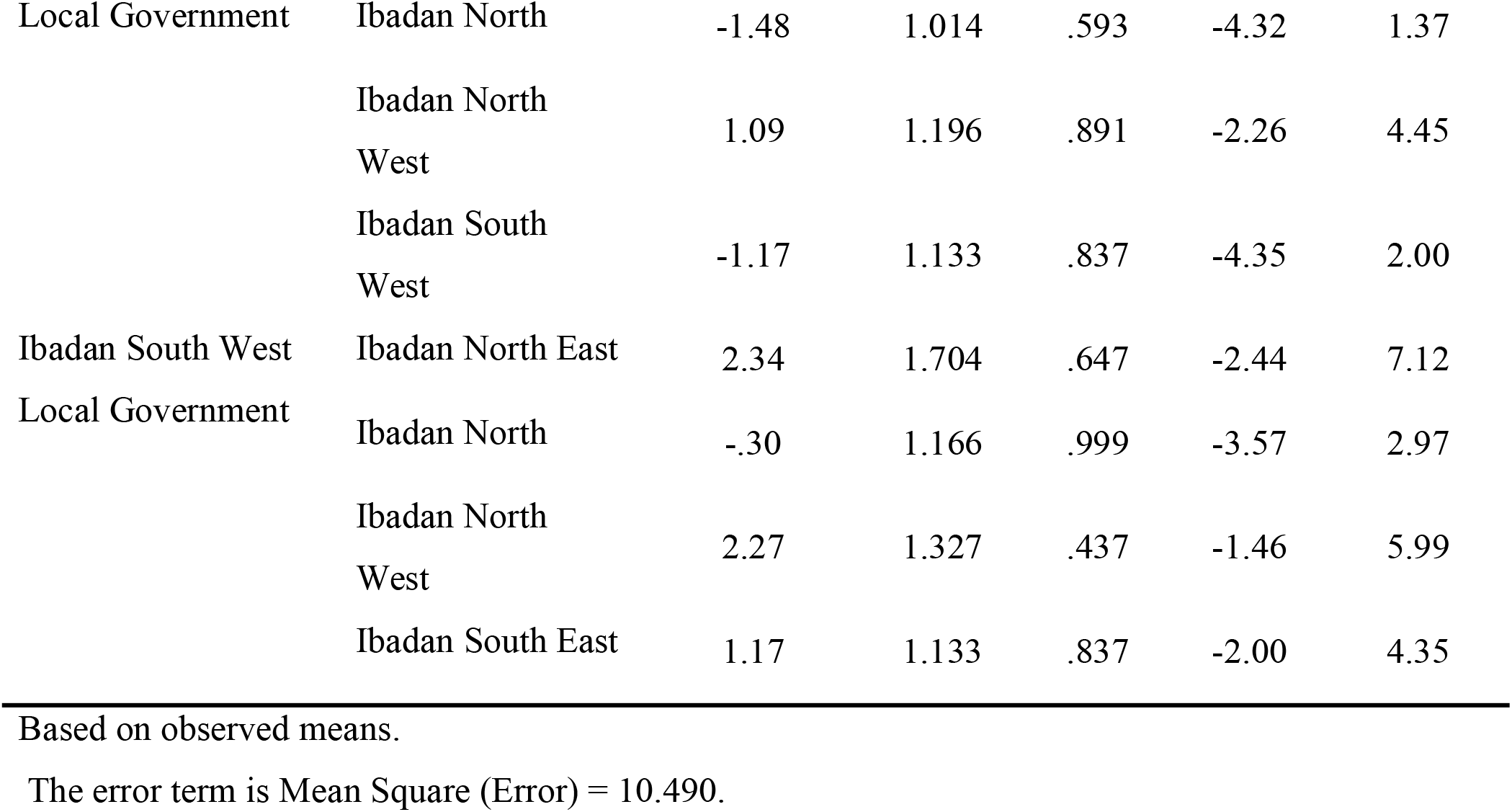
Post Hoc Test on Bird Species Richness.

The Kuskal-Wallis H Test revealed that the mean rank for Ibadan North East is **24.70**, Ibadan North is **44.74**, Ibadan North West is **25.32**, Ibadan South East is **34.55** and Ibadan South West is **36.38 (**See table 1b). It implies that mean of Ibadan North will be rank first in bird species richness, followed by Ibadan South West, Ibadan South East, Ibadan North West and Ibadan North East respectively.

The test statistics table showed a chi-square value **(*X***^**2**^**) of 8.253** and a p-value of **0.083**. At 5% C.I., the calculated p-value (0.083) is greater than 0.05. This implies that bird species richness between the different local governments is ns.

### DETERMINATION OF SPECIES ABUNDANCE WITHIN THE METROPOLIS

The results in table 4.2a showed that ***the mean bird abundance and standard deviation*** was estimated at 64.40 ± 12.28 for Ibadan North East, 68.42 ± 31.39 for Ibadan North, 46.36 ± 9.63 for Ibadan North West, 53.68 ± 24.16 for Ibadan South East and 65.69 ± 51.26 Ibadan South West. Overall mean bird abundance and standard deviation is estimated at 59.52 ± 31.35.

**Table 2a:** Descriptive Statistics on species abundance.

| **Local Government** | **N** | **Mean** | **Std. Deviation** |
| --- | --- | --- | --- |
| Ibadan North East | 5 | 64.4000 | 12.28007 |
| Ibadan North | 19 | 68.4211 | 31.39058 |
| Ibadan North West | 11 | 46.3636 | 9.63611 |
| Ibadan South East | 22 | 53.6818 | 24.16291 |
| Ibadan South West | 13 | 65.6923 | 51.26465 |
| Total | 70 | 59.5286 | 31.35571 |

**Table 2b:**
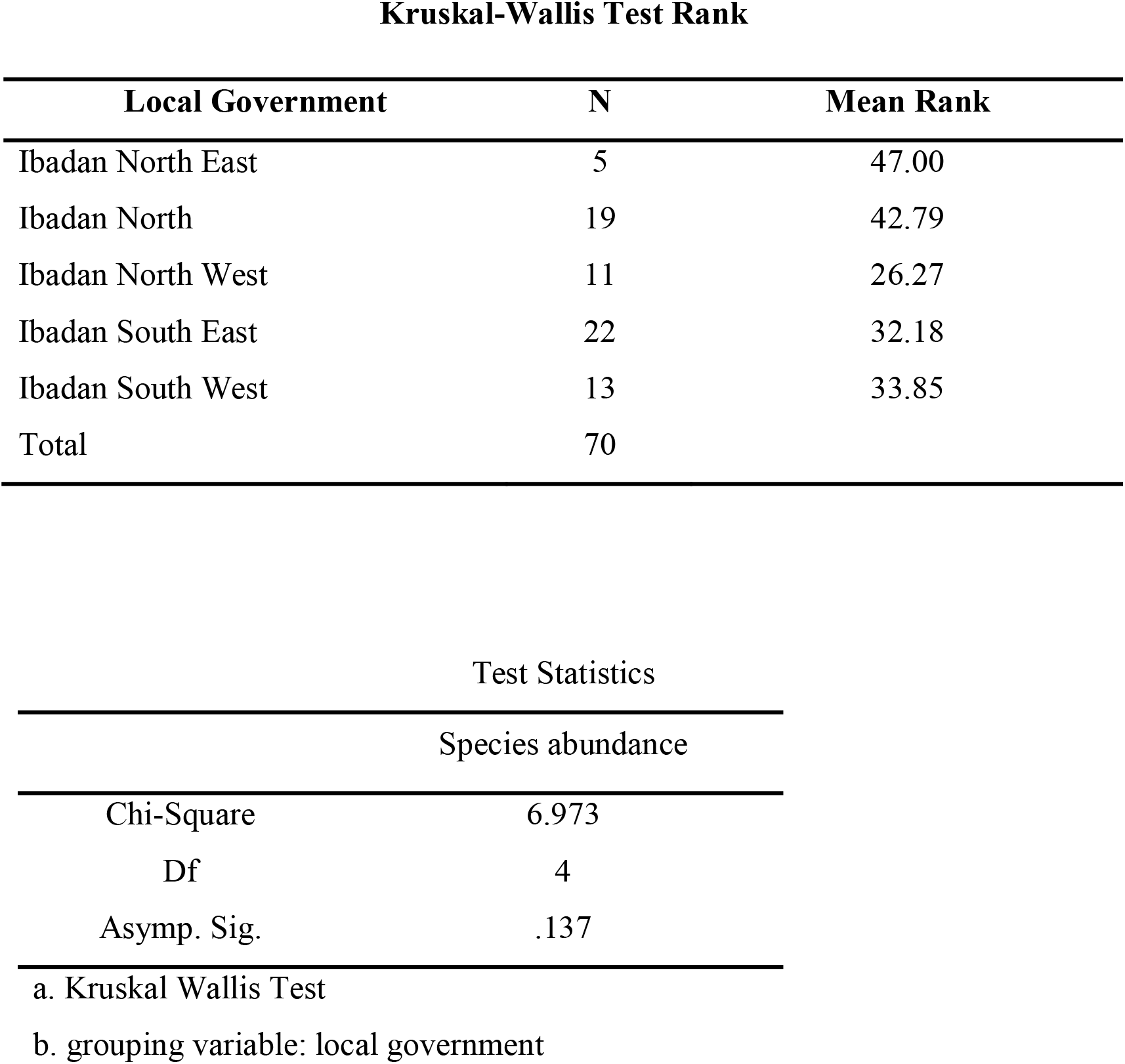
Kruskal-Wallis H Test on Bird Species Abundance.

| **Local Government** | **N** | **Mean Rank** |
| --- | --- | --- |
| Ibadan North East | 5 | 47.00 |
| Ibadan North | 19 | 42.79 |
| Ibadan North West | 11 | 26.27 |
| Ibadan South East | 22 | 32.18 |
| Ibadan South West | 13 | 33.85 |
| Total | 70 |  |

**Test Statistics**
| **Species abundance** |  |
| --- | --- |
| Chi-Square | 6.973 |
| Df | 4 |
| Asymp. Sig. | .137 |

a. Kruskal Wallis Test
b. grouping variable: local government

**Table 2c:**
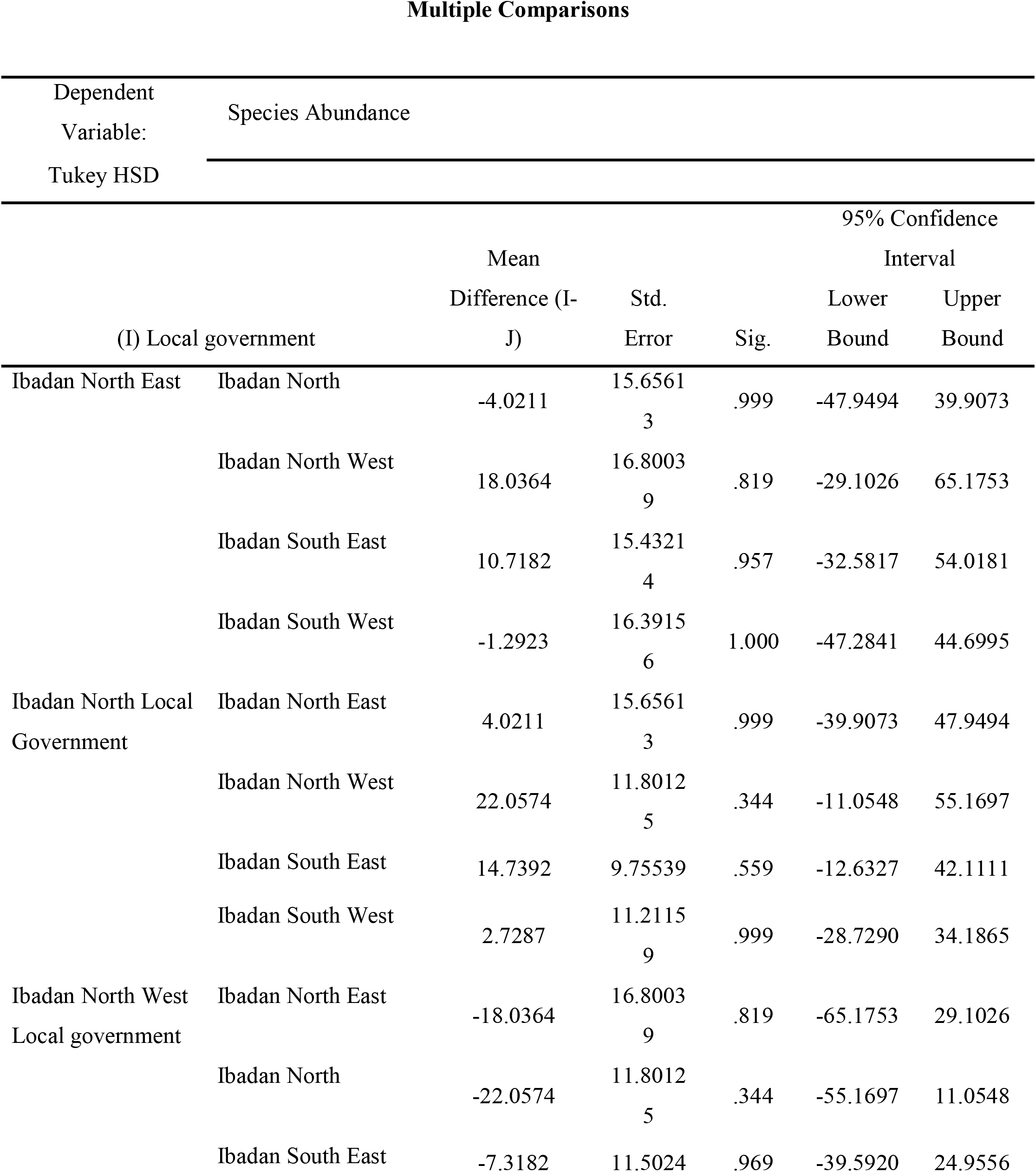
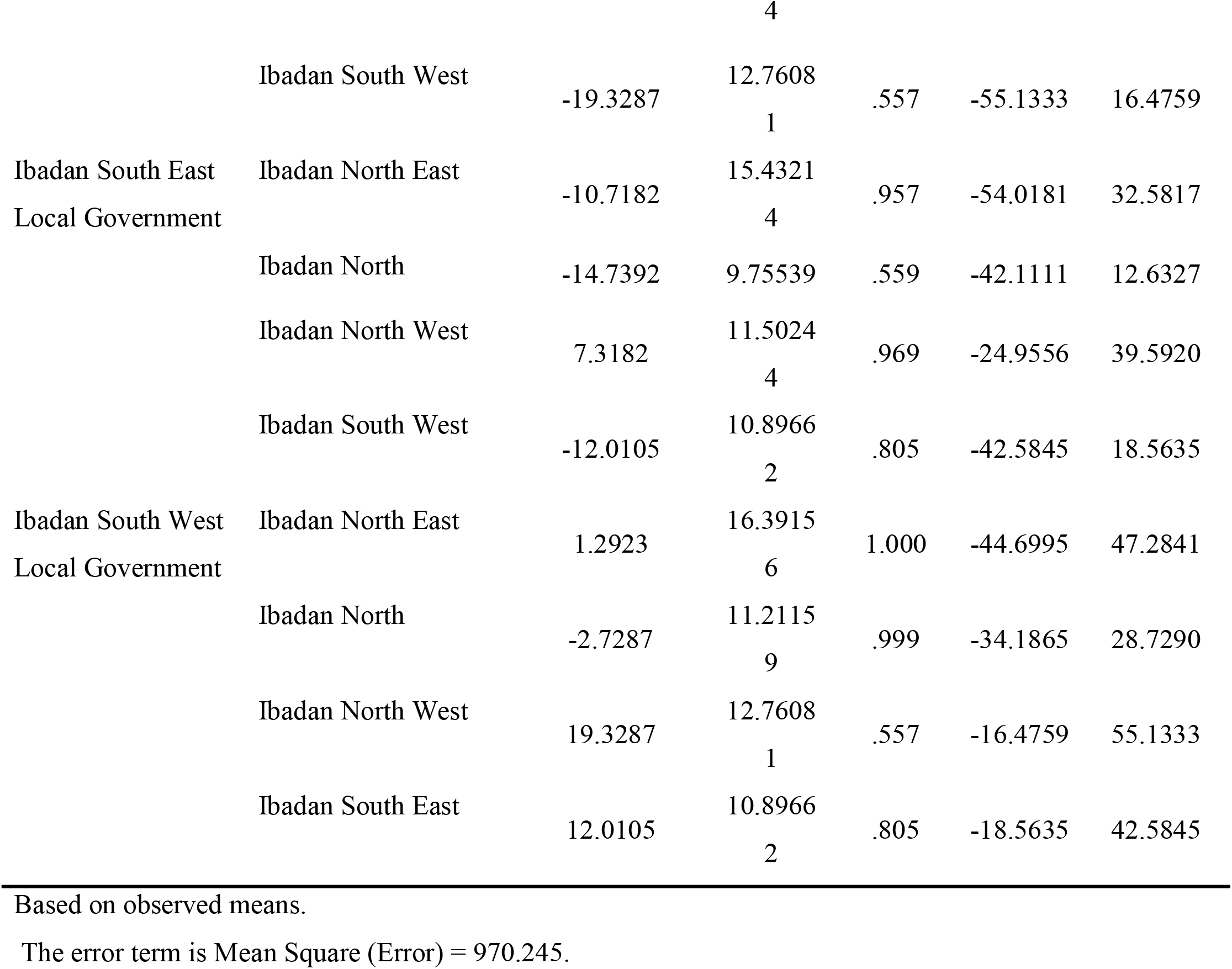
Post Hoc Test on Bird Species Abundance.

The Kruskal-Wallis Test showed that in term of ranking the abundance of bird species, Ibadan North East will be rank first, followed by Ibadan North, Ibadan North west, Ibadan South east and Ibadan South West respectively. (See table 2b)

The test statistics table showed a chi-square value **(X**^**2**^**) of 6.973** and a p-value of **0.137**. At 5% S.L., the calculated p-value is .137; which is greater than 0.05. This implies that bird species abundance between the different local governments is ns.

The multiple comparisons table showed that were no statistically significant differences in bird species abundance between any of the selected local government, as p-value in all the comparison are greater than 5%. (See table 2c)

### DETERMINATION OF HABITAT PARAMETERS THAT INFLUENCE BIRD SPECIES RICHNESS WITHIN IBADAN METROPOLIS

The test of between-subjects effects in table 3 showed whether the habitat variables (independent variables) are statistically significant in assessing and influencing bird species richness within Ibadan Metropolis.

**Table 3:** Test of between-Subjects Effects on Species Richness and habitat variables Test of Between-Subjects Effects.

| Source | Type III Sum<br>of Squares | df | Mean Square | F | Sig. (p-value) |
| --- | --- | --- | --- | --- | --- |
| Corrected Model | 247.093<sup>a</sup> | 15 | 16.473 | 1.762 | .066 |
| Intercept | 317.361 | 1 | 317.361 | 33.952 | .000 |
| Number of Buildings | .698 | 1 | .698 | .075 | .786 |
| Number of Trees | 4.202 | 1 | 4.202 | .450 | .505 |
| Number of Masts | 3.020 | 1 | 3.020 | .323 | .572 |
| Number of Paved | 7.556 | 1 | 7.556 | .808 | .373 |
| Number of Lawns | .838 | 1 | .838 | .090 | .766 |
| Percentage of Canopy<br>cover | 3.362 | 1 | 3.362 | .360 | .551 |
| Percentage of Ground<br>cover | .405 | 1 | .405 | .043 | .836 |
| Presence of farmland | .280 | 1 | .280 | .030 | .863 |
| Number of vehicles | 69.511 | 1 | 69.511 | 7.437 | .009 |
| Number of Pedestrians | 25.651 | 1 | 25.651 | 2.744 | .103 |
| Number of nests | 4.831 | 1 | 4.831 | .517 | .475 |
| Error | 504.750 | 54 | 9.347 |  |  |
| Total | 7377.000 | 70 |  |  |  |
| Corrected Total | 751.843 | 69 |  |  |  |

226 R Squared = .329 (Adjusted R Squared = .142)

The results revealed that there was **no** statistically significant relationship between the effect of the number of **buildings** on bird species richness, **F (1, 54) = 0.075, p = 0.786;** there was **no** statistically significant relationship between the effects of the number of **trees** on bird species richness, **F (1, 54) = 0.450, p = 0.505;** there was **no** statistically significant relationship between the effects of the number of **masts** on bird species richness, **F (1, 54) = 0.323, p = 0.572;** there was **no** statistically significant relationship between the effects of the number of **paved roads** on bird species richness, **F (1, 54) = 0.808, p = 0.373;** there was **no** statistically significant relationship between the effects of the number **lawns** on bird species richness, **F (1, 54) = 0.090, p = 0.766;** there was **no** statistically significant relationship between the effects of the percentage of **canopy covers** on bird species richness, **F (1, 54) = 0.360, p = 0.881;** there was **no** statistically significant relationship between the effects of the percentage of **ground covers** on bird species richness, **F (1, 54) = 0.043 p = 0.836;** there was **no** statistically significant relationship between the effects of the presence of **farm land** on bird species richness, **F (1, 54) = 0.30, p = 0.863;** there was **no** statistically significant relationship between the effects of the number of **vehicles** on bird species richness, **F (1, 54) = 7.437, p = 0.09;** there was **no** statistically significant relationship between the effects of the number of **pedestrians** on bird species richness, **F (1, 54) = 2.744, p = 0.103;** there was **no** statistically significant relationship between the effects of the number of **nests** on bird species richness, **F (1, 54) = 0.517, p = 0.475**.

Also, **R square of 32.9%** implies that all the independent variables are not too strong in predicting or influencing model.

We can therefore conclude that there were no statistically significant interaction between all the habitat parameters/variables and bird species richness within Ibadan Metropolis. That is all the habitat parameters/ variables did not influence bird species richness within Ibadan.

### DETERMINATION OF HABITAT PARAMETERS THAT INFLUENCE BIRD SPECIES ABUNDANCE WITHIN IBADAN METROPOLIS

The results revealed that there was **no** statistically significant interaction between the effect of the number of **buildings** on bird species richness, **F (1, 54) = 0.20, p = 0.888;** there was **no** statistically significant interaction between the effects of the number of **trees** on bird species richness, **F (1, 54) = 2.181, p = 0.145;** there was **no** statistically significant interaction between the effects of the number of **masts** on bird species richness, **F (1, 54) = 0.359, p = 0.552;** there was statistically significant interaction between the effects of the number **of paved roads** on bird species richness, **F (1, 54) = 5.632, p = 0.021;** there was **no** statistically significant interaction between the effects of the number of **lawns** on bird species richness, **F (1, 54) = 1.408, p = 0.241;** there was **no** statistically significant interaction between the effects of the percentage of **canopy covers** on bird species richness, **F (1, 54) = 1.412 p = 0.240;** there was **no** statistically significant interaction between the effects of the percentage of **ground cover** on bird species richness, **F (1, 54) = 0.01, p = 0.981;** there was **no** statistically significant interaction between the effects of the presence of **farm land** on bird species richness, **F (1, 54) = 0.100, p = 0.754;** there was statistically significant interaction between the effects of the number of **vehicles** on bird species richness, **F (1, 54) = 5.042, p = 0.022;** there was **no** statistically significant interaction between the effects of the number of **pedestrians** on bird species richness, **F (1, 54) = 0.302, p = 0.585;** there was **no** statistically significant interaction between the effects of the number of **nests** on bird species richness, **F (1, 54) = 0.419, p = 0.520**. The results revealed that there were statistically significant interaction between the number of paved and the number of vehicles on the bird species abundance and no statistically significant interaction in all other habitat variables/parameters. (See table 4)

**Table 4:** Test of between-Subjects Effects on Species Abundance and Habitat Variables.

| Source | Type III Sum of Squares | df | Mean Square | F | Sig. ( p - value) |
| --- | --- | --- | --- | --- | --- |
| Corrected Model | 17805.715 | 15 | 1187.048 | 1.281 | .246 |
| Intercept | 5930.968 | 1 | 5930.968 | 6.401 | .014 |
| Number of Buildings | 18.417 | 1 | 18.417 | .020 | .888 |
| Number of Trees | 2027.731 | 1 | 2027.731 | 2.188 | .145 |
| Number of Masks | 332.432 | 1 | 332.432 | .359 | .552 |
| Number of Paved | 5218.020 | 1 | 5218.020 | 5.632 | .021 |
| Number of Lawns | 1304.756 | 1 | 1304.756 | 1.408 | .241 |
| Number of Canopy cover | 1308.209 | 1 | 1308.209 | 1.412 | .240 |
| Number of Ground cover | .531 | 1 | .531 | .001 | .981 |
| Presence of farmland | 92.273 | 1 | 92.273 | .100 | .754 |
| Number of vehicles | 5135.375 | 1 | 5135.375 | 5.542 | .022 |
| Number of Pedestrians | 279.845 | 1 | 279.845 | .302 | .585 |
| Number of nests | 388.402 | 1 | 388.402 | .419 | .520 |
| Error | 50033.728 | 54 | 926.551 |  |  |
| Total | 315895.000 | 70 |  |  |  |
| Corrected Total | 67839.443 | 69 |  |  |  |

R Squared = .262 (Adjusted R Squared = .058)

Also, R square of 26.2% implies that the independent variables are weak in predicting or influencing bird species abundance.

We can therefore conclude that only number of paved roads and number of vehicles exact a significant effect on bird species abundance while all other habitat variables do not exact significant influence on bird species abundance within Ibadan Metropolis. That is, all other habitat parameters/ variables did not influence bird species abundance, except numbers of paved roads and vehicles available within Ibadan metropolis.

### 4.5 PICTURES OF BIRDS SPECIES AND ACTIVITY SIGHTED IN URBAN AREAS

**Figure 4:**
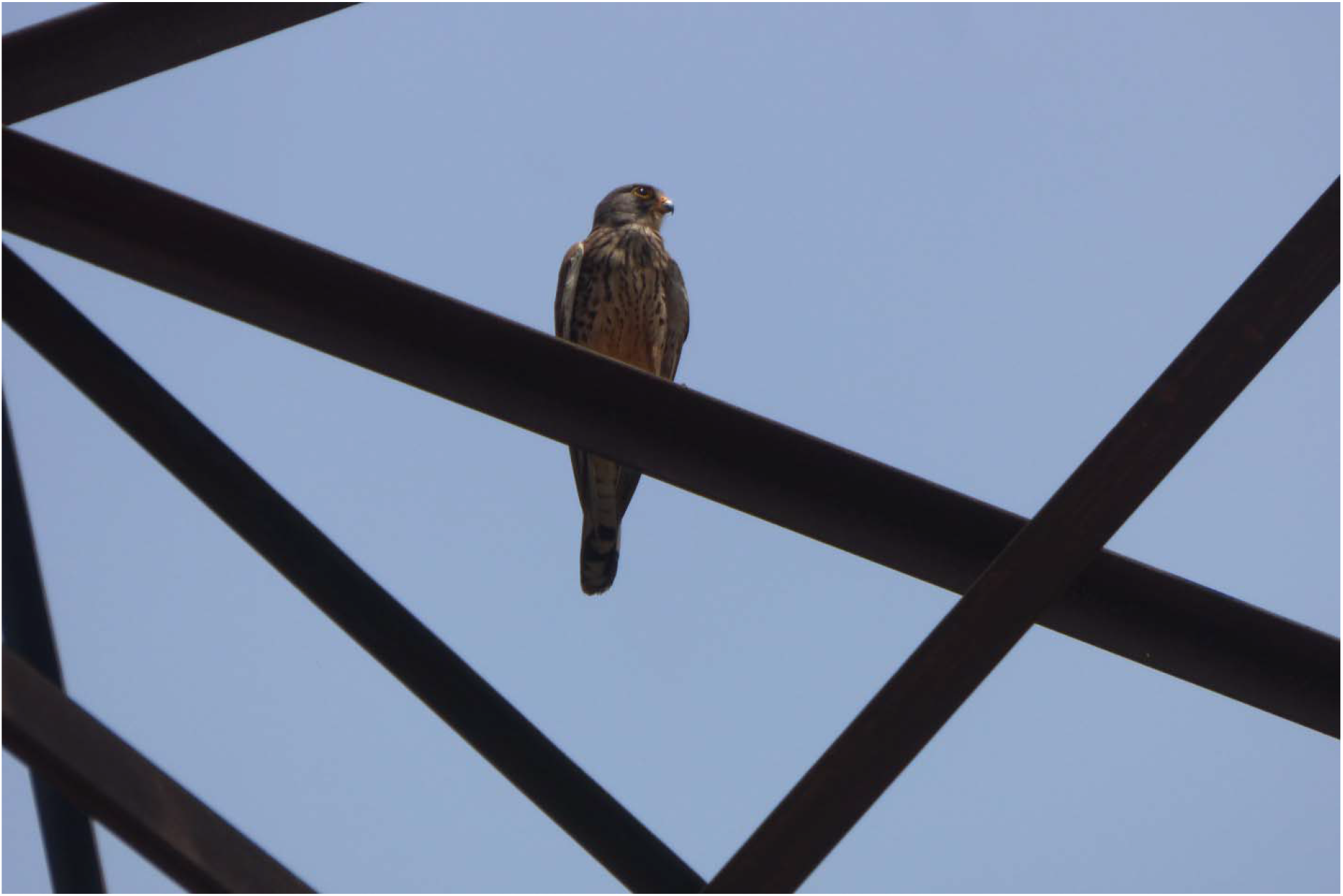
Common Kestrel (*Falco tinnunculus)* resting on a billboard

**Figure 5:**
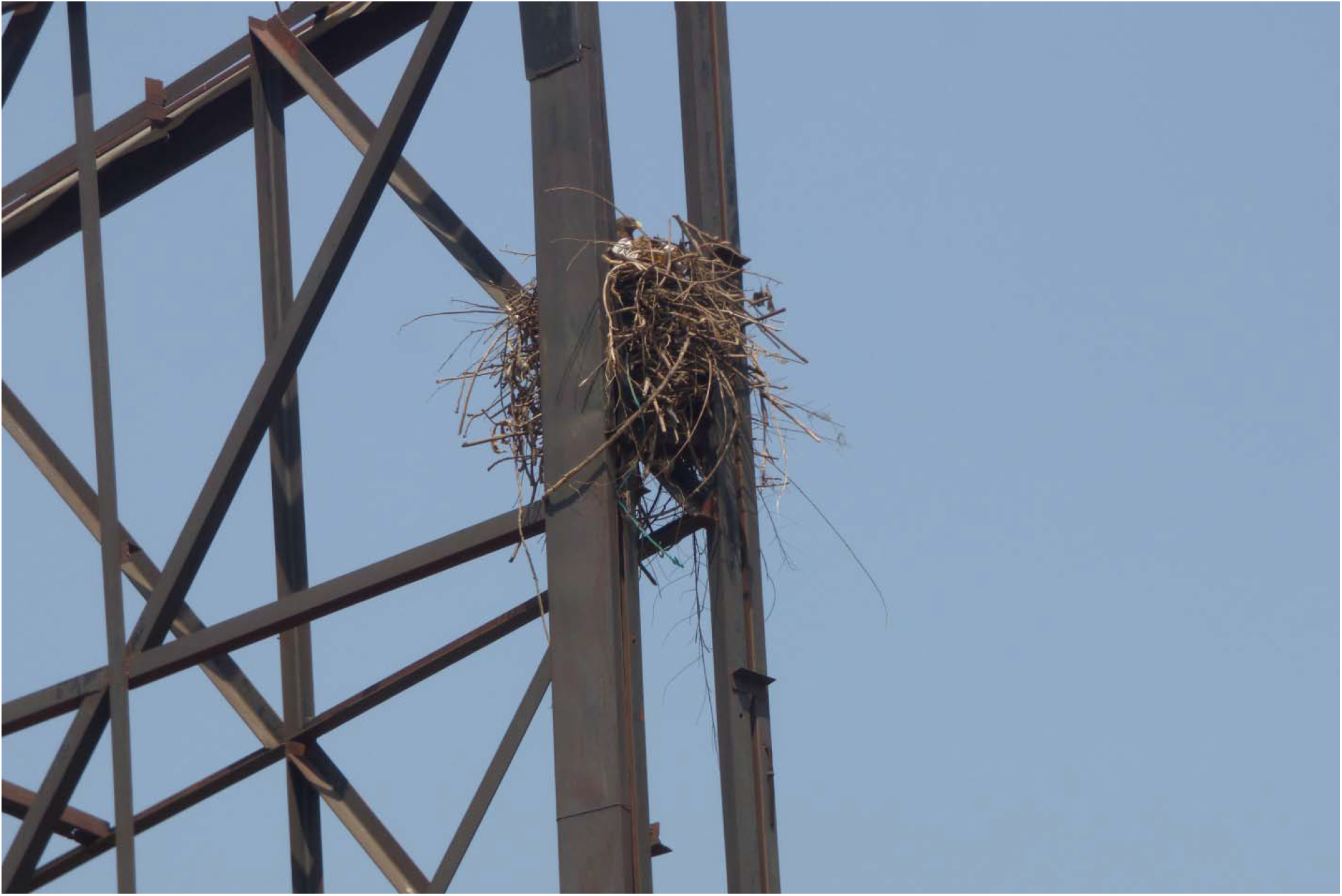
Yellow Billed Kite (*Milvus aegyptius*) brooding in a nest hung on a billboard

**Figure 6:**
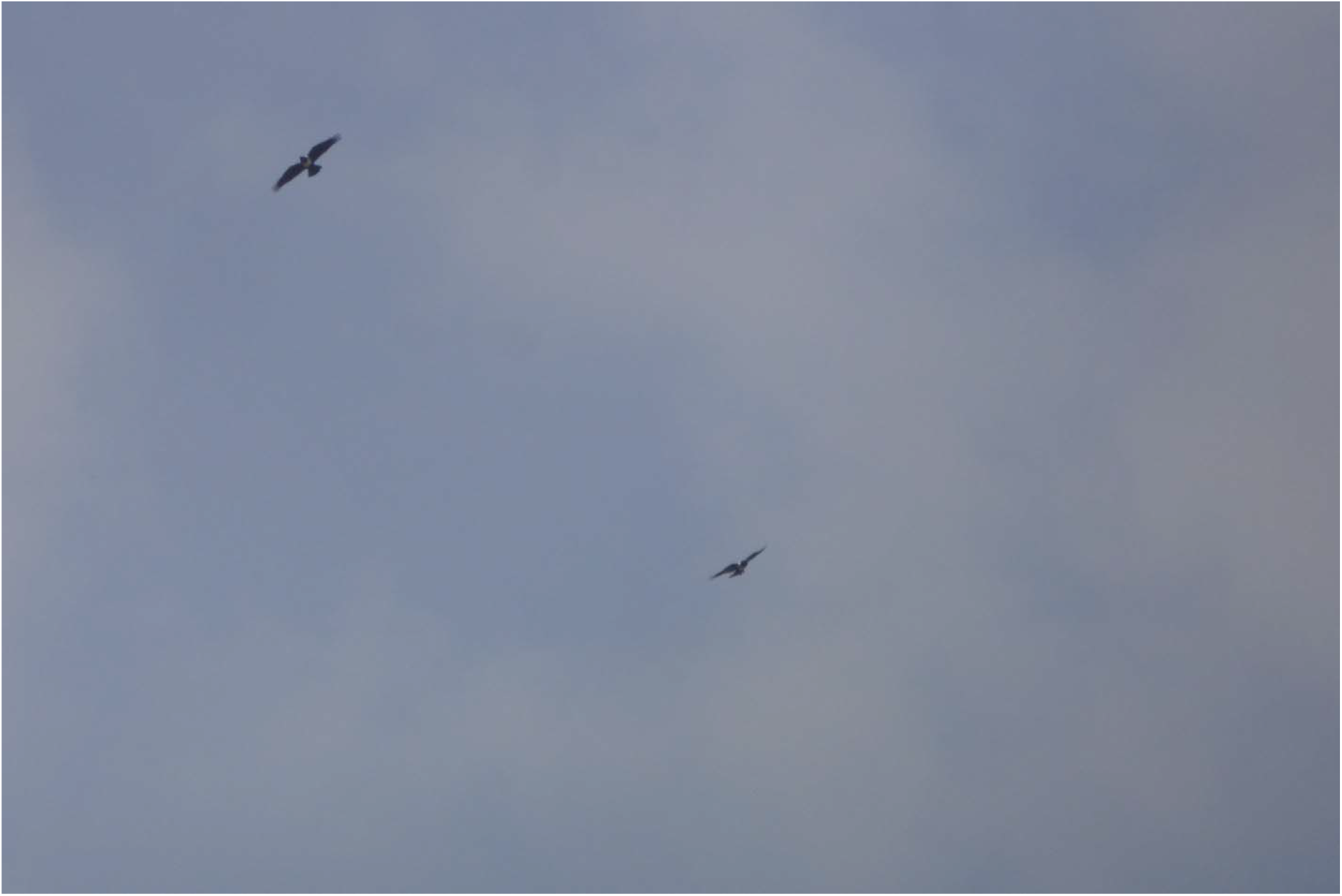
Pied Crow (*Corvus albus)* hovering in the sky

**Figure 7:**
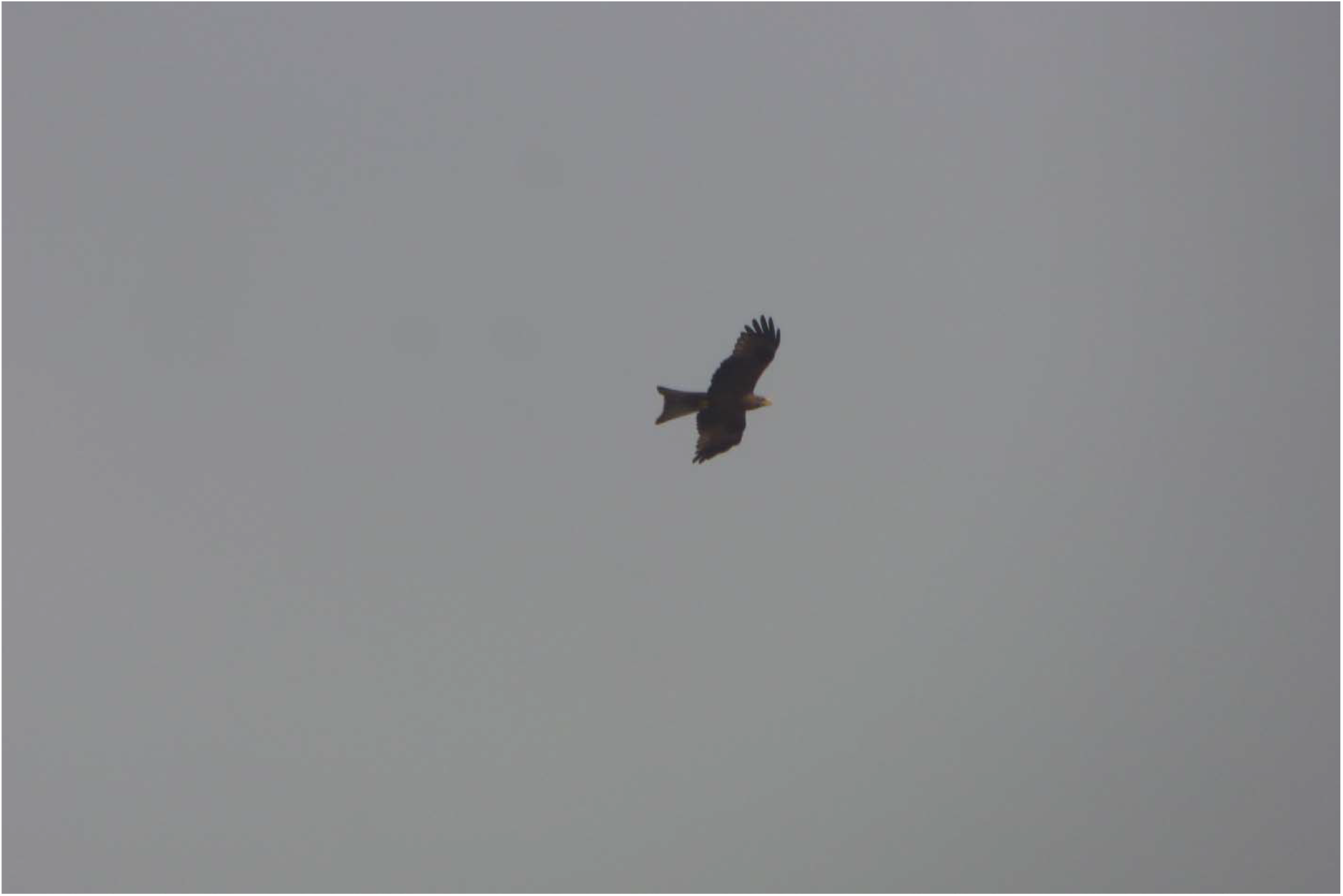
Yellow Billed Kite (*Milvus aegyptius*) hovering in the sky

**Figure 8:**
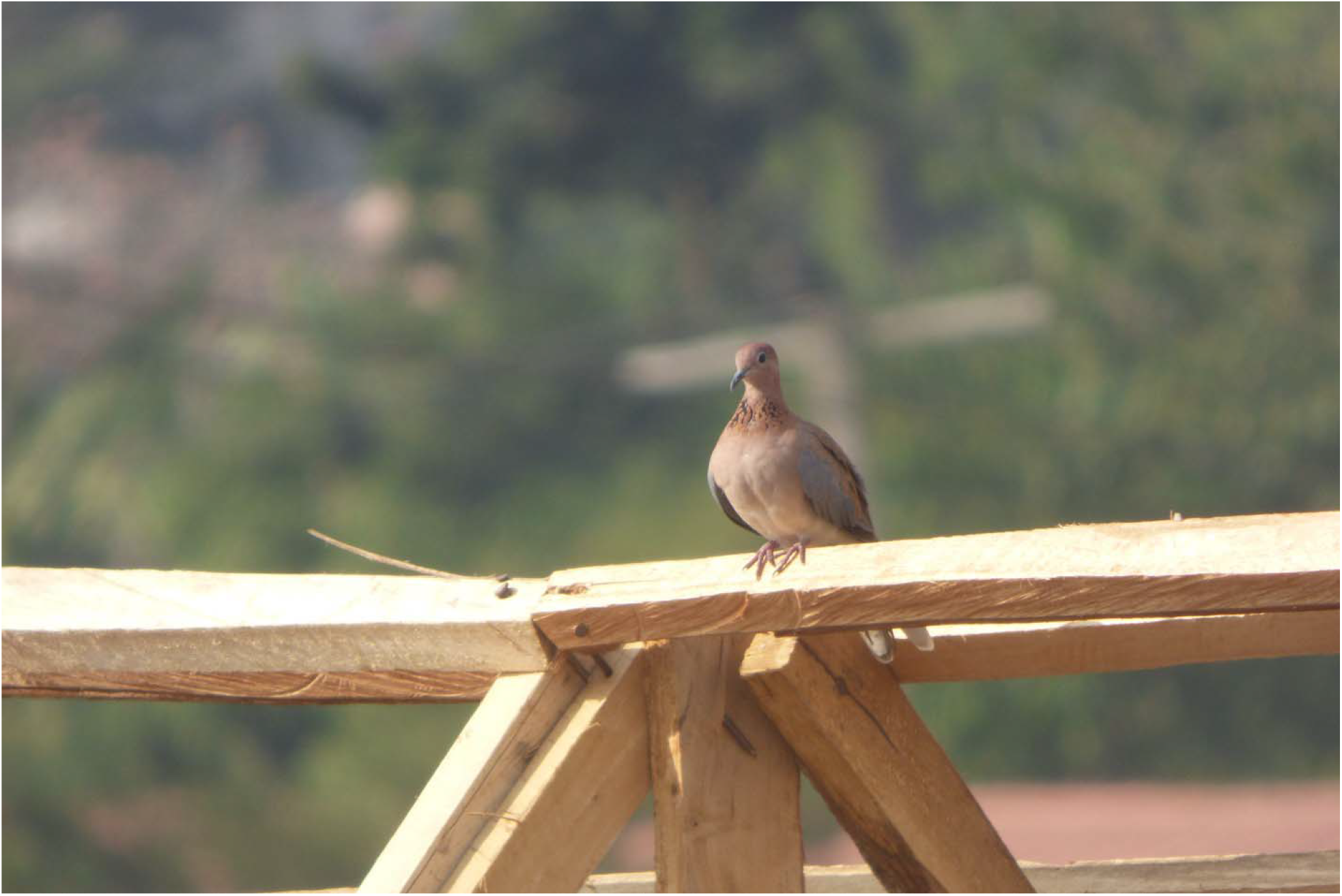
Laughing Dove (*Streptopelia semitorquata)* resting on a roof

**Figure 9:**
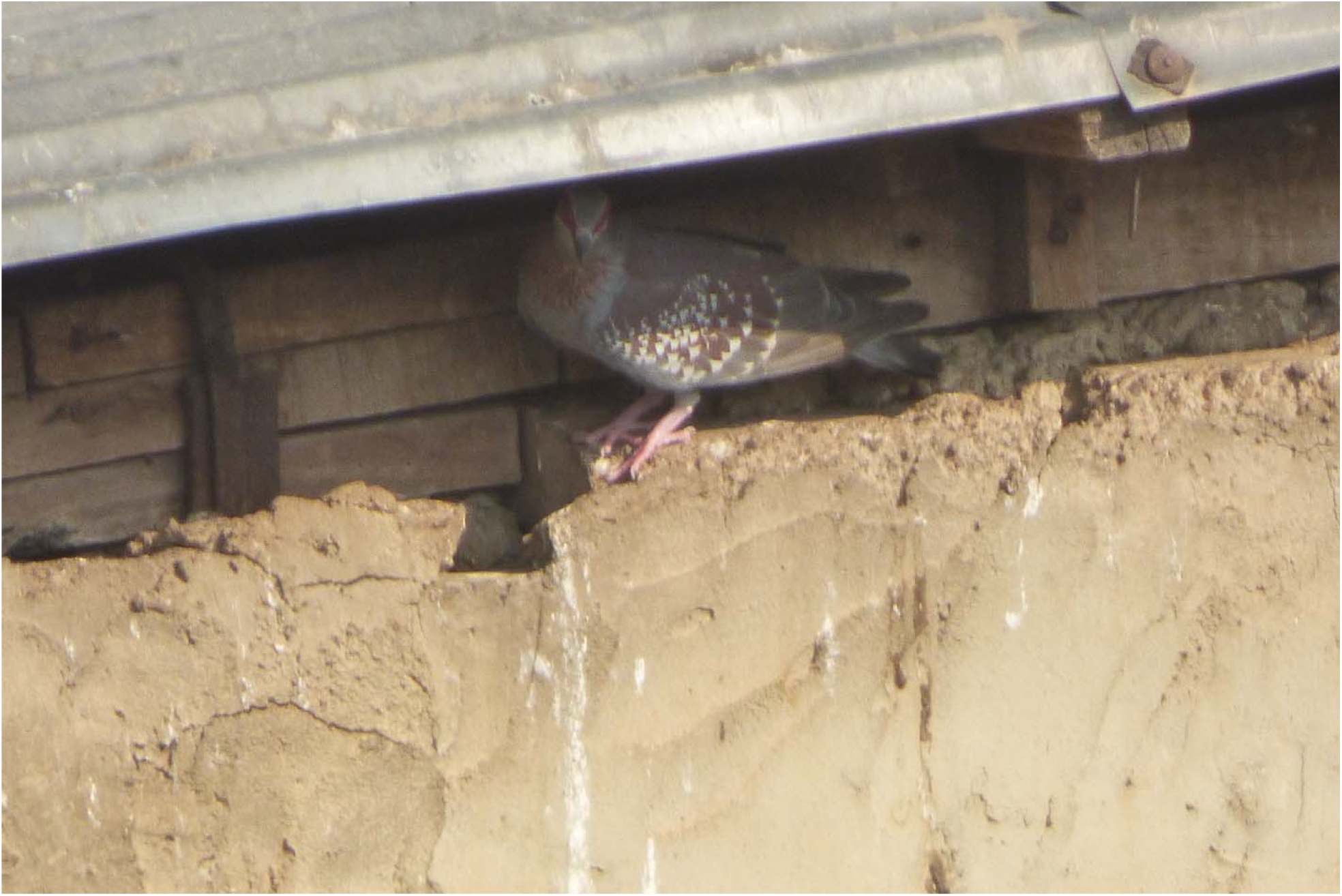
Speckled Pigeon (*Columba guinea)* in its roost

## DISCUSSION

The objective of this study was to examine the impacts of urbanization on bird species richness, abundance and the influence of habitat parameters on species richness and abundance in five selected urban local governments within Ibadan Metropolis. The researchers assessed the presence of birds using QGIS software application, of a uniform grid of 500 by 500 metres installed in some strategic areas in those local governments.

A total number of fifty-six (56) different species of birds were observed at the end of the assessment and those species were classified into Thirty (30) different family. Speckled Pigeon (*Columba guinea)* and Pied Crow (*Corvus albus*) had the highest number of individuals sighted, which were mostly found the roost around buildings’ roof. The high numbers of *C*.*albus* might be as a result of the availability of refuse dumps littering the local governments visited. *C*.*albus* can be categorized as urban exploiters. These species have adapted to exploiting urban areas as seen in their close association with human habitation and dependence on human food subsidies (Labiran and Iwajomo 2018). All bird species were recorded within the surveyed grids. Few individuals of the endangered Grey Parrot *Psittacus erithacus* (BirdLife International 2019) were recorded during this survey. All other species recorded are currently categorized as Least Concerned under the IUCN Red List (IUCN 2018).

The Kruskal-Wallis H Test on the bird species richness revealed that the mean of Ibadan North Local Government has the highest and was rank first, subsequently followed by Ibadan South West, Ibadan South East, Ibadan North West and Ibadan North East Local Government respectively. The Post Hoc Test also reveal that there were no statistically significant differences in bird species richness in any of the Local Governments. This implies bird species richness does not vary in size of availability from one local government to another. Our study revealed that there were no difference in species richness which might probably due to the fact that all the selected local governments are urban centers and virtually the same ways of life are being practiced as they are all found in the same metropolis.

The Kruskal-Wahills H Test on bird species Abundance within the Local Government revealed that the mean of Ibadan North East has the highest mean rank, followed by Ibadan North, Ibadan South West, Ibadan South East and Ibadan North West Local Governments respectively. The test of statistics showed that there was no statistically significant difference in bird species abundance between the local governments as a whole. The post Hoc Test showed there were no statistical significant difference between each unit of the local government. This implies bird species abundance does not varies or different in size of availability from one local government to another.

The Test of Between-subjects Effects was carried out to assess the influence of habitat variables on bird species richness and bird species abundance within the metropolis. There was no statistical significant effects/interaction between all the habitat variables and species richness. This implies that habitat variables do not influence bird species richness in Ibadan Metropolis. Our observed results were different compared to Iwajomo et al. (2018), where bird species richness was significantly related to the percentage of ground cover and densities of shrubs and buildings in the study area.

Furthermore, the number of Paved roads and number of Vehicles exacted a significant effect on bird species abundance while others variables under consideration did not exact statistically significant effects on bird species abundance. Generally, bird abundance has been reported to increase in response to increase in urbanization (Tietze and Arise 2018) and this increase has been attributed to the availability of food subsidies and the reduction of predation pressure (Luck and Smallbone 2010).

Thus, this study serves as a baseline to foster future research in how bird diversity is affected by urban ecosystem in Nigeria. Since urban landscapes represent a mosaic of habitats providing diverse opportunities for birds, planning efforts should seek to create and maintain an appropriate balance of habitats that provide the most opportunities for the most species. Also, for successful urban bird conservation, there is need to address the conservation needs of birds, habitat potential of various urban landscape forms, and the needs and motivations of urban residents (Labiran and Iwajomo 2018).

## ACKNOWLEDGEMENT

I give all the glory, honor and adoration to the Almighty God who has proved His greatness and support to me beyond measures during and after this research work.

My sincere appreciation goes Prof. I. A. Ayodele for his fatherly advice and concern throughout the period of this research. Profound gratitude also goes to Dr. A. T. Adeyanju for his love, relentless effort and contributions towards the successful completion of this project.

My heartfelt gratitude also goes to my ever loving and supportive parents, Mr and Mrs Adegbola, for their spiritual, moral and financial contributions. I cannot forget my paternal aunt and her husband, Surveyor and Mrs Morawo for their immense support and love. Special regards also goes to Rev. Prof. R.S Babatunde and his family, whom have been a source of support and motivation for me, thanks for the prayers and contributions.

I cannot but give due praises and appreciation to my sisters Abike Oluwadamilola Adegbola, and Oluwamayowa Susan Adegbola, for their numerous contributions at several occasions throughout my stay in the University, may the good Lord bless you more and more. I give special thanks to my cousin and her husband-Engr Bolaji and Mrs Adeola Sadik for their numerous contributions. Oyebamiji Ayokuleyin Uthman, thank you for assisting me to make this work a successful one, I am forever indebted to you. I also acknowledge Azeez Grace Jesuferanmi for her contributions to this research work even at her inconvenience.

To my course mates, the FANTABULOUS CLASS OF 40, you all are the best, I wish us all greater heights and achievements in our future endeavors. The journey would not have been fun-filled and interesting without you all. Thank you for all the good and ugly times we shared, your respect and support really means a lot. I have learnt a lot and I am very grateful. You all will always be remembered.

I also appreciate the following people: Mrs Temidayo Adeyanju, Mr Michael (Love Doctor) and Ibrahim Abdullahi and everybody who have been instrumental in making this project a success.

Thank you all and God bless you.

## Appendix SPECIES OF BIRDS OBSERVED AND THEIR GROUPING

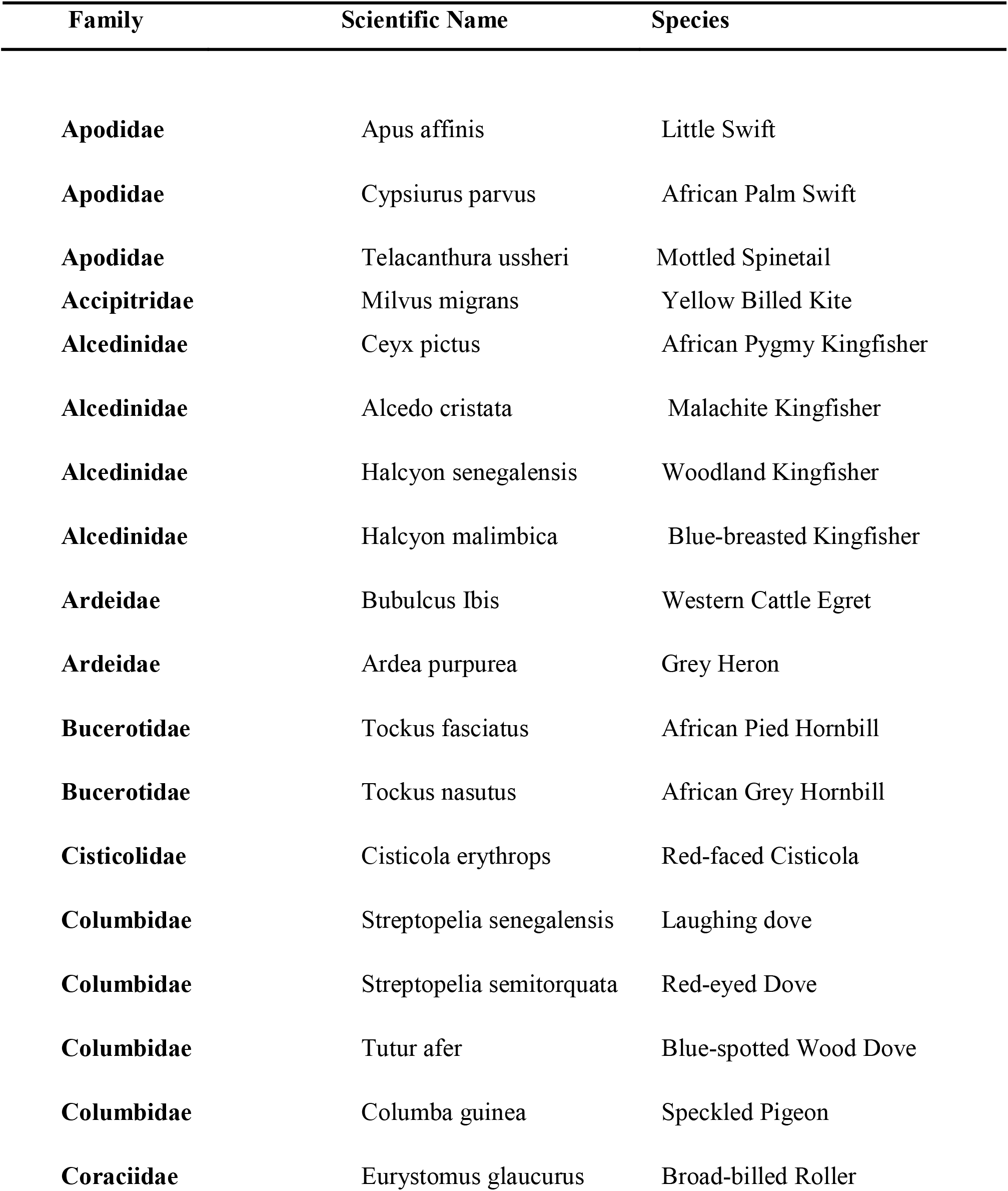
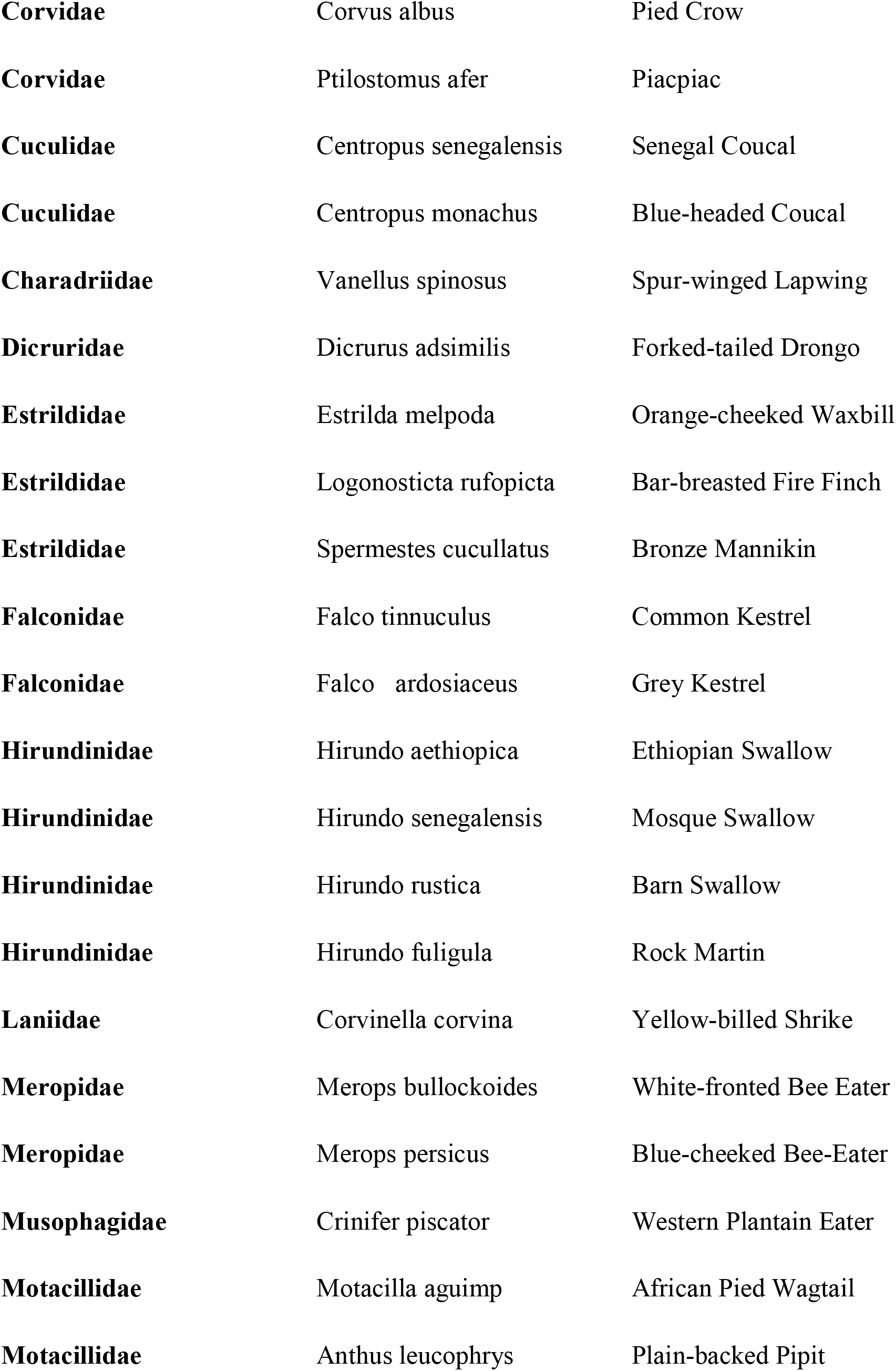
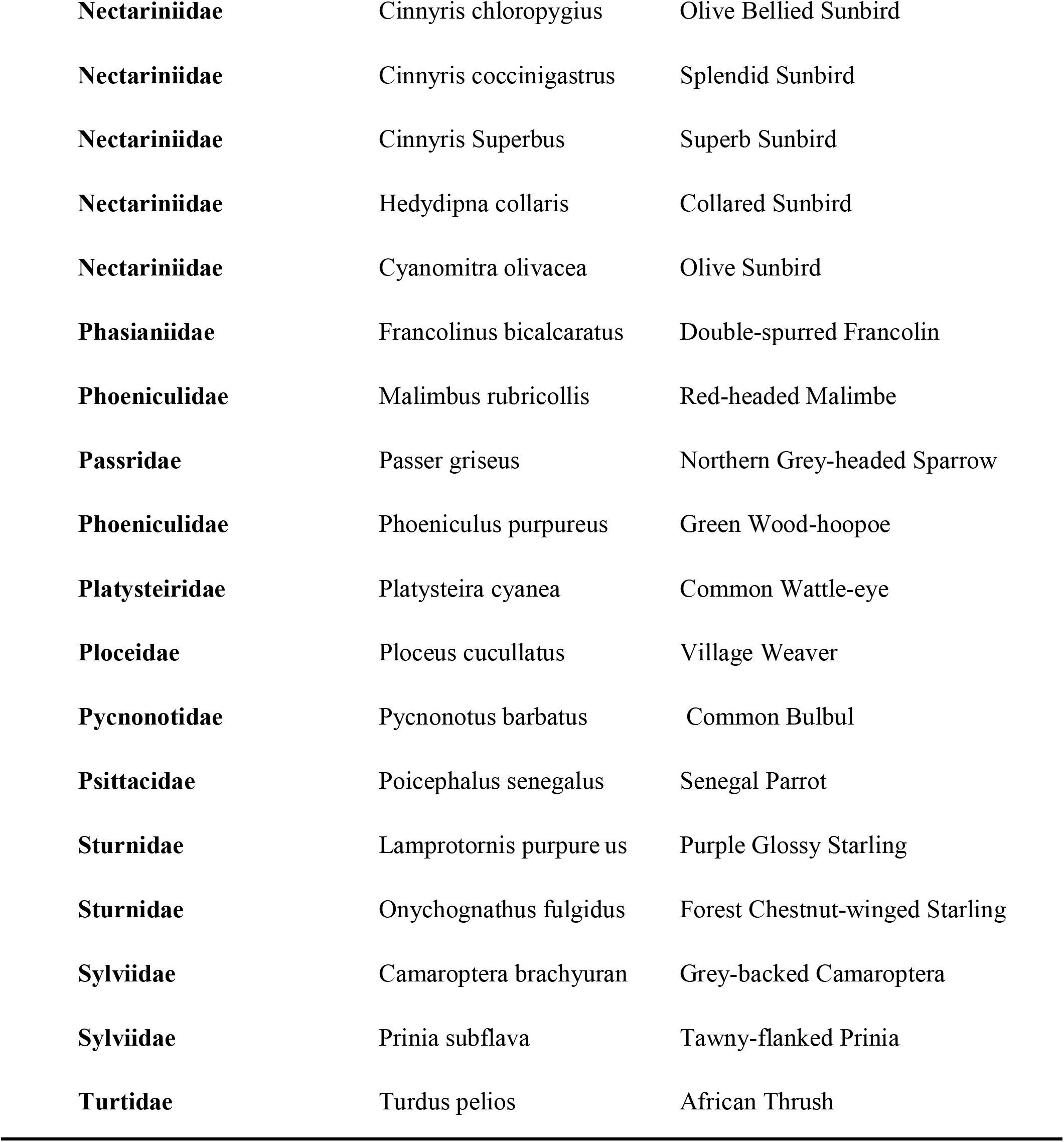

**Appendix****424 SPECIES OF BIRDS OBSERVED AND THEIR GROUPING**
|  | **Family** | **Scientific Name** | **Species** |
| --- | --- | --- | --- |
| 425 |  |  |  |
| 426 | **Apodidae** | *Apus affinis* | Little Swift |
| 427 | **Apodidae** | *Cypsiurus parvus* | African Palm Swift |
| 428 | **Apodidae** | *Telacanthura ussheri* | Mottled Spinetail |
| 429 | **Accipitridae** | *Milvus migrans* | Yellow Billed Kite |
| 430 | **Alcedinidae** | *Ceyx pictus* | African Pygmy Kingfisher |
| 431 | **Alcedinidae** | *Alcedo cristata* | Malachite Kingfisher |
| 432 | **Alcedinidae** | *Halcyon senegalensis* | Woodland Kingfisher |
| 433 | **Alcedinidae** | *Halcyon malimbica* | Blue-breasted Kingfisher |
| 434 | **Ardeidae** | *Bubulcus Ibis* | Western Cattle Egret |
| 435 | **Ardeidae** | *Ardea purpurea* | Grey Heron |
| 436 | **Bucerotidae** | *Tockus fasciatus* | African Pied Hornbill |
| 437 | **Bucerotidae** | *Tockus nasutus* | African Grey Hornbill |
| 438 | **Cisticolidae** | *Cisticola erythrops* | Red-faced Cisticola |
| 439 | **Columbidae** | *Streptopelia senegalensis* | Laughing dove |
| 440 | **Columbidae** | *Streptopelia semitorquata* | Red-eyed Dove |
| 441 | **Columbidae** | *Tutur afer* | Blue-spotted Wood Dove |
| 442 | **Columbidae** | *Columba guinea* | Speckled Pigeon |
| 443 | **Coraciidae** | *Eurystomus glaucurus* | Broad-billed Roller |
| 444 | **Corvidae** | *Corvus albus* | Pied Crow |
| 445 | **Corvidae** | *Ptilostomus afer* | Piacpiac |
| 446 | **Cuculidae** | *Centropus senegalensis* | Senegal Coucal |
| 447 | **Cuculidae** | *Centropus monachus* | Blue-headed Coucal |
| 448 | **Charadriidae** | *Vanellus spinosus* | Spur-winged Lapwing |
| 449 | **Dicruridae** | *Dicrurus adsimilis* | Forked-tailed Drongo |
| 450 | **Estrildidae** | *Estrilda melpoda* | Orange-cheeked Waxbill |
| 451 | **Estrildidae** | *Logonosticta rufopicta* | Bar-breasted Fire Finch |
| 452 | **Estrildidae** | *Spermestes cucullatus* | Bronze Mannikin |
| 453 | **Falconidae** | *Falco tinnuculus* | Common Kestrel |
| 454 | **Falconidae** | *Falco ardosiaceus* | Grey Kestrel |
| 455 | **Hirundinidae** | *Hirundo aethiopica* | Ethiopian Swallow |
| 456 | **Hirundinidae** | *Hirundo senegalensis* | Mosque Swallow |
| 457 | **Hirundinidae** | *Hirundo rustica* | Barn Swallow |
| 458 | **Hirundinidae** | *Hirundo fuligula* | Rock Martin |
| 459 | **Laniidae** | *Corvinella corvina* | Yellow-billed Shrike |
| 460 | **Meropidae** | *Merops bullockoides* | White-fronted Bee Eater |
| 461 | **Meropidae** | *Merops persicus* | Blue-cheeked Bee-Eater |
| 462 | **Musophagidae** | *Crinifer piscator* | Western Plantain Eater |
| 463 | **Motacillidae** | *Motacilla aguimp* | African Pied Wagtail |
| 464 | **Motacillidae** | *Anthus leucophrys* | Plain-backed Pipit |
| 465 | **Nectariniidae** | *Cinnyris chloropygius* | Olive Bellied Sunbird |
| 466 | **Nectariniidae** | *Cinnyris coccinigastrus* | Splendid Sunbird |
| 467 | **Nectariniidae** | *Cinnyris Superbus* | Superb Sunbird |
| 468 | **Nectariniidae** | *Hedydipna collaris* | Collared Sunbird |
| 469 | **Nectariniidae** | *Cyanomitra olivacea* | Olive Sunbird |
| 470 | **Phasianidae** | *Francolinus bicalcaratus* | Double-spurred Francolin |
| 471 | **Phoeniculidae** | *Malimbus rubricollis* | Red-headed Malimbe |
| 472 | **Passridae** | *Passer griseus* | Northern Grey-headed Sparrow |
| 473 | **Phoeniculidae** | *Phoeniculus purpureus* | Green Wood-hoopoe |
| 474 | **Platysteiridae** | *Platysteira cyanea* | Common Wattle-eye |
| 475 | **Ploceidae** | *Ploceus cucullatus* | Village Weaver |
| 476 | **Pycnonotidae** | *Pycnonotus barbatus* | Common Bulbul |
| 477 | **Psittacidae** | *Poicephalus senegalus* | Senegal Parrot |
| 478 | **Sturnidae** | *Lamprotornis purpure us* | Purple Glossy Starling |
| 479 | **Sturnidae** | *Onychognathus fulgidus* | Forest Chestnut-winged Starling |
| 480 | **Sylviidae** | *Camaroptera brachyuran* | Grey-backed Camaroptera |
| 481 | **Sylviidae** | *Prinia subflava* | Tawny-flanked Prinia |
| 482 | **Turtidae** | *Turdus pelios* | African Thrush |

